# Cortical geometry constrains the unimodal anchors of sensory integration

**DOI:** 10.64898/2026.06.13.732038

**Authors:** Alexander Holmes, Wei Wei, R. Austin Benn, Francesco Alberti, Robert Scholz, James C. Pang, Alex Fornito, Peter A. Robinson, Daniel S. Margulies

## Abstract

The human cerebral cortex is organized along a unimodal-to-transmodal hierarchy, which provides a putative substrate for the integration of sensory signals from primary cortical fields with information from other modalities. Diverse structural and functional properties of the cortex, including myelination, gene expression, neurodevelopmental timing, and inter-regional functional connectivity are patterned along this hierarchy. One exception to this hierarchy are the dominant eigenmodes of cortical geometry, which are instead patterned along rostrocaudal, mediolateral, and dorsoventral axes, each anchored by a primary sensory area at one extreme. Recent work has reconstructed the unimodal-to-transmodal hierarchy by integrating seed-based functional connectivity from three primary sensory areas, suggesting hierarchical organization may be driven by converging sensory input. Although geometric eigenmodes do not directly express the unimodal-to-transmodal hierarchy, they may encode modality-specific sensory organization originating from primary areas. Using MRI data from the Human Connectome Project, we tested whether geometric eigenmodes encode sensory function by modelling multisensory integration directly from cortical geometry. Specifically, we substituted functional connectivity maps from each primary sensory area with the rostrocaudal, mediolateral, and dorsoventral geometric eigenmodes, following a previously validated sensory integration mapping framework. Each geometric eigenmode corresponded to a distinct sensory domain (rostrocaudal-visual: |r| = 0.516; mediolateral-somatosensory: |r| = 0.551; dorsoventral-auditory: |r| = 0.342). Together, the three geometric eigenmodes created a mapping space that differentiated between unimodal brain regions with similar accuracy as fMRI-based models (δ = 64.74°; p < .001); however, differences between geometric and functional maps were largest within the transmodal association cortex. Reproducing the full unimodal-to-transmodal hierarchy required additional higher-frequency geometric eigenmodes (15 eigenmodes: r^2^ = 0.64). These findings suggest that the unimodal anchors of sensory integration are shaped by cortical geometry, with low-frequency geometric eigenmodes providing a structural basis for sensory organization, while the transmodal apex requires additional structural information to emerge.

## Introduction

A dominant feature of human cortical organization is that many molecular, cellular, physiological and anatomical properties are arranged along a spatial axis that extends from unimodal to transmodal regions (Hong et al., 2020; Huntenburg et al., 2018; Margulies et al., 2016). This axis can be mapped non-invasively in living humans functional magnetic resonance imaging (fMRI) via a modal decomposition of pair-wise functional connectivity matrices measured with resting-state fMRI (Margulies et al., 2016). The resulting modes are commonly termed *gradients* or *functional modes* (Margulies et al., 2016). The unimodal-to-transmodal axis consistently emerges as the dominant functional mode and closely aligns with classical models of hierarchical processing in the brain, whereby distinct streams of unimodal information converge within higher-order transmodal regions (Jung et al., 2022; Mesulam, 1998, 2012; Smallwood et al., 2021). Consistent with this view, recent work has found that linear combinations of seed-based functional connectivity originating from visual, somatosensory, and auditory seed regions captures hierarchical organization as multisensory stimuli across domains converges within the default mode network (Wei et al., 2024).

Beyond fMRI, numerous studies have shown that many different properties of the cortex are spatially patterned along the unimodal-to-transmodal axis, including regional myelination (Grydeland et al., 2019; Paquola et al., 2019), and cortical folding (Hill, Dierker, et al., 2010; Li et al., 2013), areal expansion through development and evolution (Hill, Inder, et al., 2010; Raznahan et al., 2011), and timing of neurodevelopmental maturation (Larsen et al., 2023; Sydnor et al., 2021, 2023). For example, primary cortical areas exhibit near-adult surface area from birth (Hill, Inder, et al., 2010), whereas higher-order transmodal regions mature progressively through infancy, childhood, and adolescence, driven largely by increasing cortical folding that supports the development of complex cognitive functions (De Vareilles et al., 2023; Gilmore et al., 2012; Lyall et al., 2015). These convergent findings position the unimodal-to-transmodal axis as a major organizing principle of cortical structure and function (Larsen et al., 2023; Sydnor et al., 2021, 2023).

An unresolved question concerns how this dominant unimodal-to-transmodal axis of cortical organization emerges. Several processes have been proposed, including the development of long-range cortico-cortical and thalamocortical connections (Park et al., 2024). However, more fundamental constraints may be imposed by the geometry of the cortex itself. For instance, Margulies et al. (2016) showed that the transmodal peaks of the unimodal-to-transmodal axis are positioned in locations that are equidistant from the primary sensory fields V1, A1, and S1. Early transcriptional gradients that pattern the cortex follow simple geometric axes, such as those extending along rostrocaudal, dorsoventral, and mediolateral directions, as do gradients of the timing of regional neurogenesis (Alfano et al., 2014; O’Leary & Nakagawa, 2002; O’Leary & Sahara, 2008). Notably, these axes correspond to the first three non-global modes, called geometric eigenmodes of the cortex, representing the dominant axes of geometric variation in the cortex (Pang et al., 2023a).

Under neural field theory, a popular class of biophysical models for large-scale brain dynamics, these modes also represent preferred, or resonant, standing wave patterns of excitation (Gabay & Robinson, 2017; Robinson et al., 2003, 2016). Consistent with this view, geometric eigenmodes can be used to parsimoniously account for diverse aspects of brain function (Pang et al., 2023a), connectivity (Normand et al., 2025), and regionalization (Pang et al., 2025).

We have recently shown that the dominant modes of cortical function in neonates more closely resemble these geometric modes and gradually transition to a unimodal-to-transmodal gradient between the ages of 3 and 8 years (Holmes et al., 2026). Such findings support the idea that the unimodal-to-transmodal hierarchy gradually emerges from an initial organization dictated by geometry (Fornito et al., 2025).

Notably, the first three non-global geometric eigenmodes appear to be anchored in primary sensory regions: the visual cortex occupies an extreme caudal position along the rostrocaudal mode, the somatosensory cortex occupies a central position by the central sulcus in the mediolateral mode, and the auditory cortex lies ventrally near the superior temporal gyrus in the dorsoventral mode (see Fig 1) (Holmes et al., 2026; Robinson & El Zghir, 2025). This observation implies that the specific location of primary sensory fields may result from simple geometric constraints on early transcriptional gradients (Cadwell et al., 2019). The gradual functional differentiation of these sensory fields may then drive the emergence of the unimodal-to-transmodal axis (Fornito et al., 2025). This view accords with recent evidence that seed-based resting state functional connectivity from V1, S1, and A1 can be used to model hierarchical integration in transmodal regions (Wei et al., 2024).

**Figure 1:**
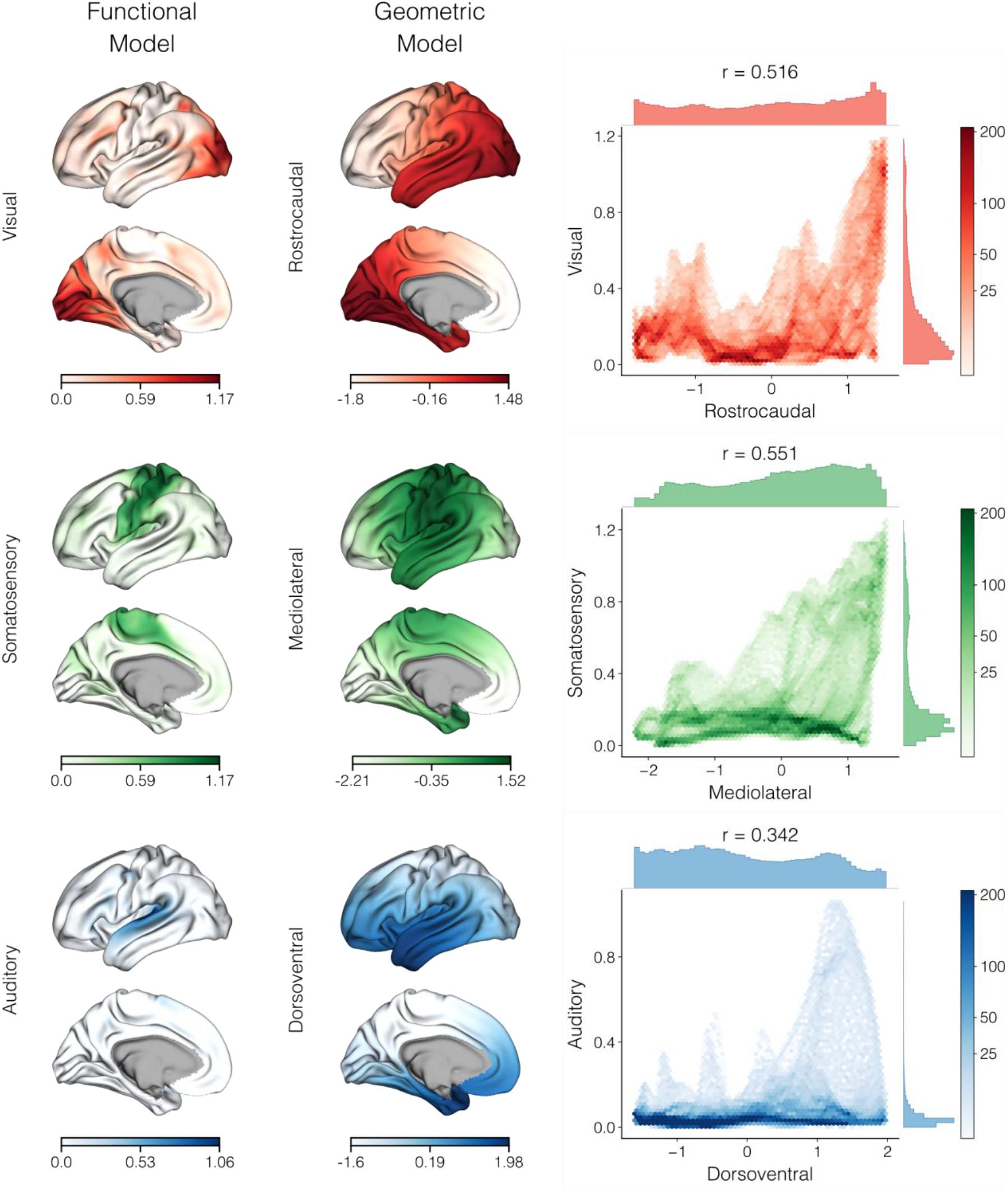
Correspondence between sensory functional connectivity and the dominant geometric eigenmodes. Left column: whole-brain functional connectivity maps from primary visual (V1), somatosensory (S1), and auditory (A1) seed regions, projected onto the cortical surface. Middle column: the first three non-constant geometric eigenmodes of the cortical surface, exhibiting rostrocaudal, mediolateral, and dorsoventral spatial patterns. Right column: vertex-wise scatter plots of each sensory connectivity map against the corresponding geometric eigenmode, with Pearson’s r reported above each panel and point density indicated by color saturation on a logarithmic scale.

We hypothesized that the eigenmodes of cortical geometry would capture the unimodal anchor points of the unimodal-to-transmodal axis, as mapped via modal decomposition of fMRI functional connectivity, which could subsequently reconstruct the hierarchical organization in the brain. We tested this assumption using a novel sensory integration mapping technique, which involved substituting beta maps of sensory activity in fMRI with the first three non-constant geometric eigenmodes, thereby estimating functional organization through a purely geometric approach.

## Methods

### Participants and MRI Preprocessing

This study used resting-state fMRI data from 167 subjects from the Human Connectome Project – Young Adult cohort (HCP-YA; mean age = 29.4 years, SD = 3.24 years; 101 females, 66 males), a publicly accessible and open-access dataset, where the sensory integration model has been applied previously (Wei et al., 2024). Scanner, acquisition, and recruitment details can be found in the original reference paper (Glasser et al., 2013).

Because the cortical sheet is a continuous, curved surface, its geometry can be represented using Laplace–Beltrami operator (LBO) eigenmodes. Thus, our analyses used a symmetric version of FreeSurfer’s fsaverage population-averaged cortical surface meshes, downsampled to 32,492 vertices in each hemisphere (Van Essen et al., 2012). We specifically derived geometric eigenmodes from the midthickness surface mesh, using the LaPy Python library (Reuter et al., 2006; Wachinger et al., 2015) to solve the eigenvalue problem for the LBO, in line with previous studies (Pang et al., 2023a). Despite using this mesh, the basic patterns of the first three non-constant geometric eigenmodes are largely invariant across different cortical surfaces and cortex-like shapes. This is because their long wavelengths render their spatial pattern insensitive to fine-scale geometric variations (Pang et al., 2023b).

### Derivation of cortical geometric eigenmodes

The geometry of the cortical surface can be modelled using the eigenmodes of the Laplace-Beltrami operator (LBO), which appears in the Helmholtz equation (Pang et al., 2023a; Robinson et al., 2016; Silberstein & Nunez, 1995). The LBO describes spatial variations on curved manifolds such as the cortical surface, with metrics relating to the lengths, angles, and curvature at each point of the manifold and is generally defined as (Chavel, 1984):

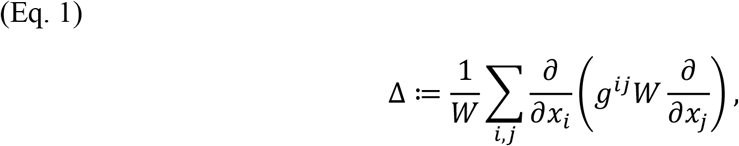

where *x*_*i*_, *x*_*j*_ are the local coordinates, *g*^*ij*^ is the inverse of the inner product metric tensor 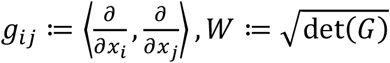, det denotes the determinant, and *G* ≔ (*g*_*ij*_).

Eigendecomposition of the LBO returns a basis set of eigenmodes and eigenvalues:

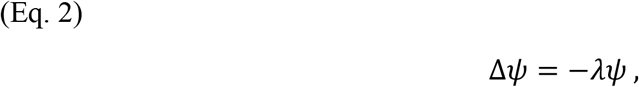

where Δ is the LBO, *ψ* is the set of eigenmodes, *ψ* = {*ψ*_1_, *ψ*_2_, … }, and eigenvalue, λ is the corresponding family of eigenvalues, λ = {λλ_1_, λλ_2_, … }. Each eigenvector corresponds to a resonant mode, or eigenmode, of the cortical geometry, which represents a specific spatial standing wave pattern that oscillates through time. The mode is resonant because it reflects a preferred way in which the cortex can respond to an input, given its geometry. The eigenmodes, *ψ*, are arranged by the corresponding eigenvalue, λ, which in turn are associated with the spatial wavelength of the modes, such that

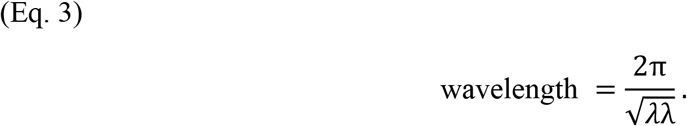

Here, increasing eigenvalues, λ, are associated with monotonically decreasing wavelengths. For example, the first eigenmode, *ψ*ψ_1_, is spatially constant, while the next three eigenmodes, *ψ*_2_, *ψ*_3_, *ψ*_4_, exhibit clear, low-frequency zones of opposing positive and negative amplitude, divided by a single nodal line (zero amplitude zones). Eigenmodes beyond this point reflect more complex patterns with higher spatial frequencies and lower wavelengths. Using a spherical approximation (Gabay & Robinson, 2017; Koussis et al., 2025; Pang et al., 2023a; Robinson et al., 2016), because the cortex is nearly topologically equivalent to a sphere, the eigenvalues and their corresponding modes of similar spatial scale and wavelength can be grouped, termed eigengroups, such that the approximate wavelength of each eigengroup corresponds to using a spherical approximation.:

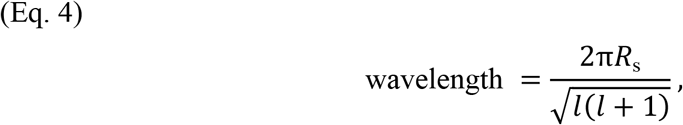

where *R*_s_ is the radius of the spherical representation of the cortex and *l* is the eigengroup number. The number of eigenmodes contained within each eigengroup increases with eigengroup number, such that each eigengroup, *l*, contains 2*l* + 1 eigenmodes and the first *l* eigengroups will contain the first *l*^2^ + 2*l* eigenmodes (when omitting the global zeroth eigenmode, which is entirely constant). The first 1000 non-constant eigenmodes span 31 eigengroups, with their specific eigengroup and wavelength reported in Table S1.

### Sensory Integration Mapping

We adapted the sensory integration mapping framework from (Wei et al., 2024) for geometric eigenmodes. Originally applied in fMRI, the model first extracts spatially averaged timeseries from three unimodal seed regions: the primary visual cortex (V1), the primary somatosensory cortex (S1), and the primary auditory cortex (A1). These regions were defined according to the HCPMMP1 parcellation (Glasser et al., 2016), which categorizes brain regions from multimodal anatomical and functional connectivity data. The sensory integration framework then aims to predict whole-brain resting-state fMRI timeseries using these three regional timeseries as predictors, represented through the following general linear model (GLM):

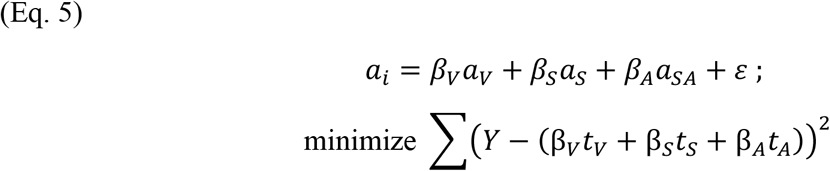

where *a*_*i*_, *a*_*V*_, *a*_*S*_, *a*_*A*_ are the timeseries of a given vertex, *i*, V1, S1, and A1, respectively. The regional beta coefficients, *β*, thereby represent the strength of functional connectivity between vertex *i* and each primary seed area in V1, S1, and A1.

The sensory functional connectivity then forms the basis for a low-dimensional polar space. Here, each primary region is mapped to one of three equidistant angles (e.g., 0°, 120°, 240°) forming the initial coordinate framework. Each modelled vertex is assigned an angular value describing its overall sensory profile. Specifically, functional preferences towards a given sensory modality are quantified based on the difference in angle from these poles, as follows:

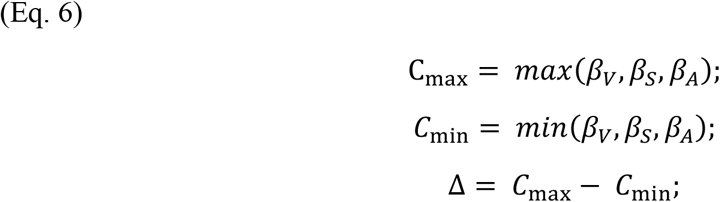

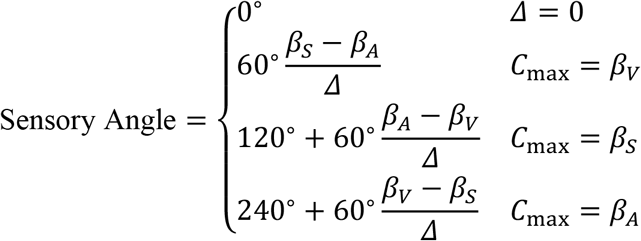

where *C*_*max*_ and *C*_*min*_ are the maximum and minimum sensory functional connectivity values, respectively, and the sensory angle is bound from 0° to 360°.

Vertices with a preferential affinity towards a single specific modality will possess sensory angles closely corresponding to that sensory pole (e.g., 0°, 120°, 240° for visual, somatosensory, and auditory seeds, respectively). In contrast, regions involved in multisensory integration possess an intermediary angle, positioned somewhere between these pre-defined poles (e.g., 60°, 180°, 300°).

Sensory angle captures relative modality preference but not the overall strength of sensory coupling. Thus, visual areas V1 and V5 may both be classified as visually dominant despite occupying different hierarchical positions (Felleman & Van Essen, 1991).

Accordingly, the sensory integration model introduces a secondary dimension, termed sensory magnitude, that identifies vertices with functional profiles that are most resemble primary areas, regardless of any specific sensory modality. Sensory magnitude is defined as the model fit of Eq. 5:

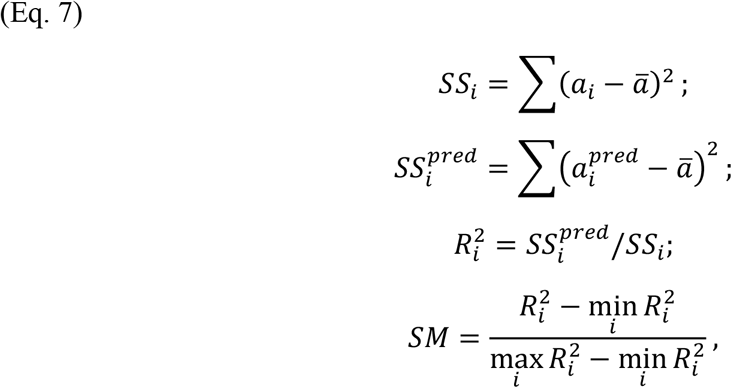

where *a*_*i*_ is the empirical timeseries at vertex *i*, 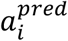 is a simulated timeseries at vertex *i* predicted from the GLM of Eq. 5, ā is the globally averaged timeseries, and *SM* is the sensory magnitude.

Along this dimension, cortical areas that are explained by primary sensory function possess higher sensory magnitude, regardless of modality. In contrast, transmodal association areas with reduced sensory functional connectivity possess lower magnitude. This unimodal-to-transmodal differentiation causes maps of whole-brain sensory magnitude to resemble the first principal gradient of functional connectivity, which adheres to a similar unimodal-to-association axis (Margulies et al., 2016; Sydnor et al., 2021, 2023; Wei et al., 2024).

Thus, sensory integration maps create a two-dimensional framework, using sensory angle to characterize multisensory function across three primary domains and sensory magnitude to contextualize this function through a hierarchical lens. A schematic workflow diagram of this method is further provided in Supplementary Materials in Fig. S1 (adapted from Wei et al., 2024 under CC BY 4.0).

### Incorporating Geometric Eigenmodes within Sensory Integration Mappings

Sensory functional connectivity is partly distance-dependent: connectivity is strongest near the corresponding primary cortex and decreases with distance (Ercsey-Ravasz et al., 2013), while transmodal peaks are maximally distant from sensory landmarks, such as the central sulcus and calcarine sulcus (Margulies et al., 2016). Because low-frequency geometric eigenmodes encode broad spatial proximity relationships on the cortical surface, they may approximate the distance-dependent component of sensory functional connectivity. We therefore tested whether geometric eigenmodes could substitute for sensory functional connectivity maps within the sensory integration framework, by replacing the sensory functional connectivity maps with the rostrocaudal, mediolateral, and dorsoventral geometric eigenmodes.

By using geometric eigenmodes in place of sensory functional connectivity maps, *β*_*V*_, *β*_*S*_, *β*_*A*_, the geometric approach skips Eq. 5 and begins with Eq. 6 to calculate the sensory angle. Accordingly, sensory magnitude was instead calculated for the geometric model by applying a GLM using geometric eigenmodes to predict the first functional gradient, which is a known proxy of the unimodal-to-transmodal hierarchy in fMRI (Margulies et al., 2016):

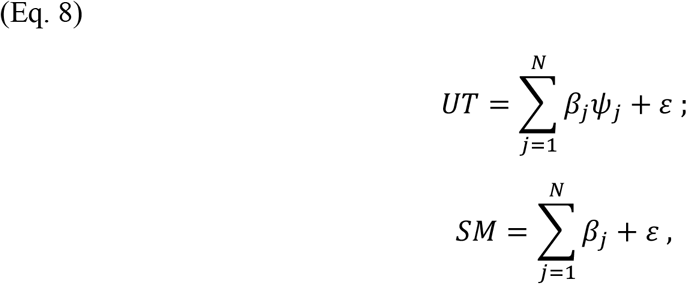

where *ψ* is the eigenmode, *N* is the number of eigenmodes included within the GLM (such as the first three eigenmodes), and *UT* is the unimodal-to-transmodal hierarchy estimated from functional connectivity data in previous work (Margulies et al., 2016).

Because sensory magnitude within the geometric model was defined relative to the functional gradient, analyses of sensory magnitude should be interpreted as representing the performance of the geometric model in reconstructing the initial reference hierarchy, rather than an independent discovery of the hierarchy.

Model accuracy was quantified as the difference in sensory angle between geometric and functional models. Smaller angular differences indicate that the geometric model is more similar to the functional reference. Angular differences were calculated at each vertex as:

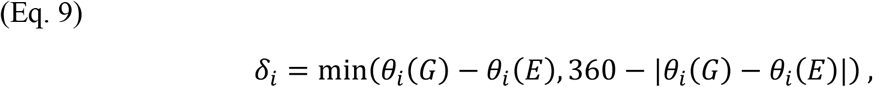

where *δ*_*i*_ is the angular difference at a given vertex *i*, and *θ*_*i*_(*G*) and *θ*_*i*_(*E*) are the sensory angles for the geometric and functional models, respectively.

We used non-parametric permutation testing via 10,000 spin tests to determine whether the observed angular differences between the geometric and functional models were smaller than expected by chance.

Finally, to assess the relationship between model accuracy and position along the unimodal-to-transmodal hierarchy, we also calculated the angular difference between models as a function of sensory magnitude. In addition to sensory magnitude, we also assessed angular differences across the 7 network labels of the Yeo-Krienen atlas (Yeo et al., 2011) and the five cortical types in the Von Economo and Koskinas cytoarchitectonic atlas (García-Cabezas et al., 2020; von Economo & Koskinas, 1925). If geometric eigenmodes uniquely align with sensory functional connectivity, then model accuracy between geometric and functional models should likely be most similar among unimodal, rather than transmodal cortical areas.

## Results

The first three non-constant geometric eigenmodes exhibited broad spatial patterns, aligned with the principal anatomical axes of the cortex (Fig. 1 left column). The first eigenmode followed a rostrocaudal axis, with extrema located near the occipital visual cortex and anterior cortical regions. The second eigenmode followed a mediolateral axis, spanning somatosensory regions toward lateral cortical territories. The third eigenmode followed a dorsoventral axis, with extrema near superior cortical regions and the temporal cortex, including auditory-related areas.

Sensory functional connectivity maps from seeded primary areas V1, S1, and A1 showed similar spatial patterns (Fig. 1 middle column). Visual functional connectivity was highest within the occipital cortex and progressively decreased with distance from primary visual areas. Somatosensory functional connectivity was highest around the central sulcus, while auditory functional connectivity was highest on the superior temporal lobe. Across modalities, the strongest values were locally clustered near the respective primary sensory cortices and diminished with distance away from these anchors.

### Correspondence between geometric eigenmodes and sensory function

Consistent with prior work (Holmes et al., 2026; Pang et al., 2023; Robinson et al., 2016), the peaks and antinodes of the dominant eigenmodes coincided with the locations of primary sensory fields, with the rostrocaudal, mediolateral, and dorsoventral modes respectively aligning with visual, somatosensory, and auditory cortical territories. Sensory functional connectivity was highly to moderately correlated with the first three geometric eigenmodes (rostrocaudal-visual: |r| = 0.516, mediolateral-somatosensory: |r| = 0.551, dorsoventral-auditory: |r| = 0.342; Fig. 1 right column). As the sign of geometric eigenmodes is arbitrary, all eigenmodes were aligned to have positive amplitude within primary areas.

However, changes in signs only influenced the direction of the correlation, not the strength.

### Estimating sensory magnitude and unimodal-to-transmodal hierarchical organization from cortical geometry

To calculate the sensory magnitude of the sensory integration model, we performed a GLM using the first three geometric eigenmodes to predict the unimodal-to-transmodal hierarchy. However, a GLM using only the three geometric eigenmodes explained limited variance in the unimodal-to-transmodal axis (r^2^ = 0.25; Fig. 2). Thus, following past work (Pang et al., 2023a), we incorporated an increasing number of geometric eigenmodes to better estimate sensory magnitude. Model performance increased to r^2^ = 0.64 with 15 eigenmodes, corresponding to the first three eigengroups. The functional model using sensory functional connectivity maps yielded a similar accuracy of r^2^ = 0.71. By 100 eigenmodes, model performance appeared to plateau at r^2^ = 0.92, with additional analyses including up to 1000 eigenmodes reported in Supplementary Materials (Fig. S2).

**Figure 2:**
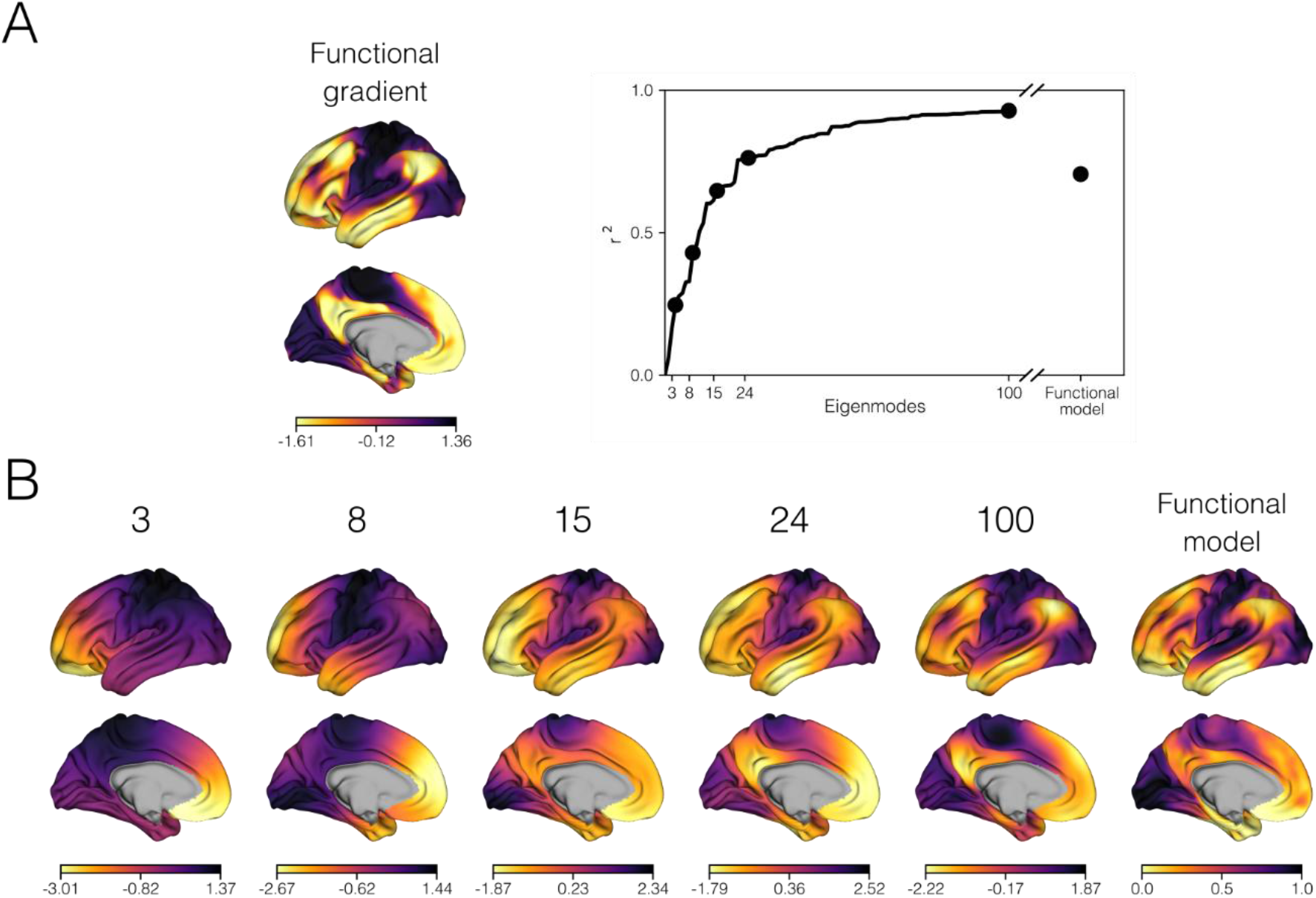
Reproduction of sensory magnitude and the unimodal-to-transmodal hierarchy from the geometric model, compared against sensory magnitude from the functional model. (A) Model r^2^ of both geometric and functional models when predicting the first functional gradient (left) (Margulies et al., 2016). Dots within the scatterplot (right) mark the model r^2^ from predictions using the first 3 eigengroups, 100 eigenmodes, and the functional model, respectively. As the first *l* eigengroups will contain a cumulative total of *l*^2^ + 2*l* eigenmodes (when omitting the constant zeroth eigenmode), the first eigengroup spans the first three eigenmodes, first and second eigengroups collectively span the first eight, and so on. (B) The sensory magnitude of the geometric model, estimated with an increasing number of geometric eigenmodes (numbered) and the sensory magnitude of the functional model. The inclusion of additional eigenmodes causes the sensory magnitude of the geometric model to more closely resemble the unimodal-to-transmodal pattern of the first functional gradient.

Because higher-order geometric eigenmodes show divergent patterns across individuals (Chen et al., 2022, 2023), we focused on a parsimonious model using the first 15-eigenmodes. This model captured major transmodal features, including the default mode network and angular gyrus, that were absent when using only 3 or 8 eigenmodes (Fig. 2).

### Sensory integration mapping through geometric eigenmodes

Having calculated sensory magnitude using 15 eigenmodes, we subsequently estimated sensory angle using only the first three eigenmodes. Alongside the geometric model, we also obtained a baseline functional model using sensory functional connectivity, previously published elsewhere (Wei et al., 2024). Qualitatively, sensory angles from the geometric model using only three eigenmodes cleanly separated primary visual, somatosensory, and auditory cortex, with these areas labelled in red, green, and blue, respectively (Fig. 3A). Notably, these patterns in sensory angle from the geometric model also resembled the functional model, which exhibited similar delineations between primary areas.

**Figure 3:**
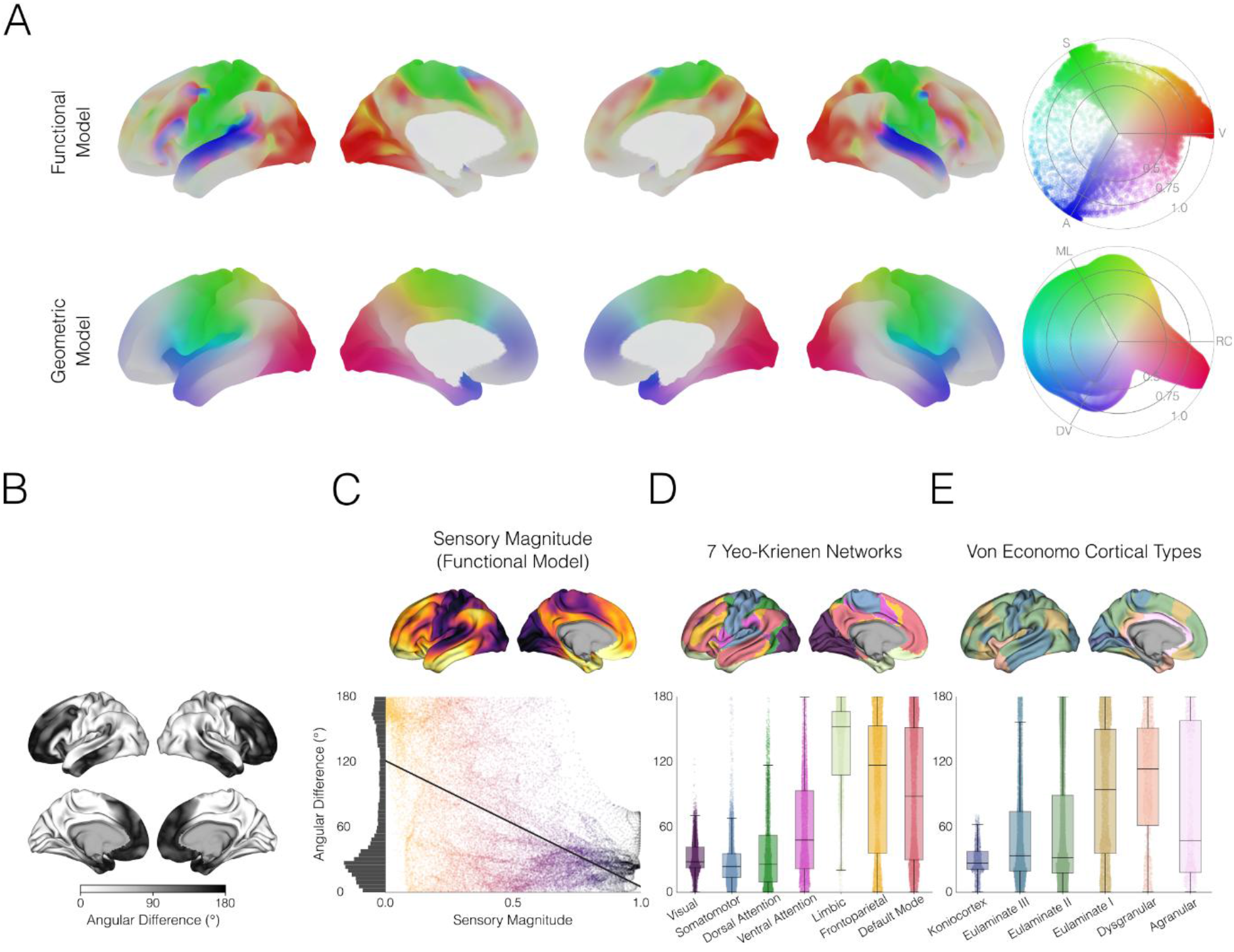
(A) Sensory integration maps from the functional model derived from resting-state fMRI data (top) and the geometric model (bottom). RGB colormaps reflect the vertex-wise sensory angle of each model, with the geometric model using the first three eigenmodes to estimate angle. The visual and rostrocaudal pole are plotted in red, the somatosensory and mediolateral pole are plotted in green, and the auditory and dorsoventral pole are plotted in blue across both models. Greyscale overlay maps reflect the sensory magnitude of each model, with the geometric model using the first 15 eigenmodes to estimate magnitude. Darker grey reflects areas with lower sensory magnitude. Across both functional and geometric models, sensory angles separated primary sensory areas V1, S1, and A1, while sensory magnitude delineated between unimodal and transmodal cortical areas. (B) Angular differences between functional and geometric models. Darker areas in the association cortex exhibited greater differences between models. (C) Scatterplot of angular difference against sensory magnitude from the functional model. The angular difference between models decreases as a function of sensory magnitude; areas with lower sensory magnitude (e.g., transmodal areas) diverged more between models than high sensory magnitude areas (e.g., primary areas). (D) Angular difference within the 7 Yeo-Krienen networks (labelled above).Visual and somatomotor networks possessed the lowest angular differences, while limbic, frontoparietal, and default mode networks possessed the greatest differences. (E) Angular difference within the five Von Economo-Koskinas cortical types (labelled above). The koniocortex possessed the lowest angular differences, while dysgranular areas possessed the greatest differences.

However, as the input parameters differed between models, the polar coordinate spaces were not innately aligned between models. Specifically, the angular poles at 0°, 120°, 240° corresponded to visual, somatosensory, and auditory functional connectivity in the functional model and rostrocaudal, mediolateral, and dorsoventral axes in the geometric model. While the input sensory functional connectivity and geometric eigenmodes were correlated (Fig. 1), some sensory angles were slightly offset between models as a result (Fig. 3A). For example, the visual cortex was centered at 0 degrees in the functional model, compared to approximately 20 degrees in the geometric model. This offset likely arises due to the differences in spatial scale between geometric eigenmodes and sensory functional connectivity. For example, the dorsoventral eigenmode captures large-scale fluctuations in whole-brain cortical geometry that extend beyond the auditory cortex.

Despite regional variations, the mean angular difference between the two models was lower than expected under spin-based null models (δ = 64.74°), suggesting that geometric eigenmodes predict sensory organization better than expected for random maps with matched spatial autocorrelation. Post-hoc permutation spin testing using 10,000 rotations demonstrated that the observed angular difference fell outside the null distribution, yielding a highly significant effect (p < .001; Fig. S3).

Most angular differences occurred in the transmodal association cortex (Fig. 3B) and there was a pronounced negative relationship between regional angular difference and sensory magnitude (Fig. 3C), suggesting that model correspondence varied with hierarchical organization. When comparing angular differences across the 7 Yeo-Krinen networks (Fig. 3D), the smallest differences were observed in visual (δ = 28.02°) and somatomotor (δ = 23.73°) networks, with the largest differences in the limbic (δ = 152.22°) networks. A similar pattern was observed across cortical types, as defined by the Von Economo and Koskinas cytoarchitectonic atlas (Fig. 3E), with lower differences in koniocortex (δ = 26.66°) and higher differences in dysgranular cortex (δ = 113.50°).

## Discussion

By substituting the functional connectivity of seeds placed in primary sensory fields with compact measures of cortical geometry, we showed that low-frequency geometric eigenmodes approximate key features of sensory integration maps. Individually, each map of sensory functional connectivity was represented by a specific eigenmode of cortical geometry, such that one of the poles of the mode co-localized with a distinct sensory region. Specifically, the rostrocaudal mode characterized visual function, the mediolateral mode characterized somatosensory function, and the dorsoventral mode characterized auditory function. Together, these three geometric eigenmodes created an angular space that differentiated unimodal sensory regions and approximately reproduced the sensory integration map. However, the differentiation of transmodal from unimodal features required additional input from successive geometric eigenmodes beyond the first three alone.

Together, these results suggest that unimodal aspects of multisensory integration are shaped by cortical geometry, while the emergence of the transmodal hierarchy requires structural information beyond the dominant geometric eigenmodes alone.

### Sensory functional connectivity maps are characterized by the dominant eigenmodes of cortical geometry

The first three non-constant geometric eigenmodes were significantly associated with sensory functional connectivity, with the strongest correlations observed between the rostrocaudal eigenmode and visual functional connectivity and the mediolateral eigenmode and somatosensory functional connectivity. Three key factors likely explain the correspondence between geometric eigenmodes and the functional organization of sensory systems:

First, the regional anchors of the geometric eigenmodes co-localize with the locations of the primary fields. The visual and somatosensory cortex occupy extreme posterior and central positions on the cortex (Brodmann, 1909), which closely match the peaks of the rostrocaudal and mediolateral eigenmodes (Pang et al., 2023; Robinson et al., 2016). As such, these eigenmodes closely correspond with their respective sensory functional connectivity maps. In contrast, the auditory cortex is located within the lateral sulcus in the upper temporal lobe (Brodmann, 1909), whereas the anchor of the dorsoventral mode is located on the ventral aspect of the temporal pole (Pang et al., 2023; Robinson et al., 2016). Accordingly, the correspondence between the auditory functional connectivity map and dorsoventral mode was weaker than those observed for vision and somatosensation.

Second, geometric eigenmodes capture broad distance-dependent properties of cortical organization. Connectivity strength generally decreases with increasing separation between cortical regions, consistent with exponential-distance rule-like relationships observed across mammalian cortex (Douglas & Martin, 2004; Ercsey-Ravasz et al., 2013; Markov et al., 2014). Because low-frequency eigenmodes encode long-wavelength spatial gradients across the cortical surface, they naturally approximate these large-scale proximity relationships.

Third, the anatomical and functional organization of sensory systems is known to be dominated by such distance-dependent factors. Sensory functional connectivity is typically strongest near the corresponding primary cortex and decreases with increasing distance from that local anchor (Ercsey-Ravasz et al., 2013). Similarly, transmodal regions, which are separated from primary sensory areas along the unimodal-to-transmodal hierarchy, are maximally distant from opposing sensory structures (Margulies et al., 2016). Because sensory processes conform to such distance-dependent properties, they are well approximated by low-frequency geometric eigenmodes, which capture broad spatial proximity relationships.

Several factors may contribute to the weaker correspondence observed for auditory functional connectivity. It is possible that the pole of the dorsoventral eigenmode may be closer to the prospective position of A1 early in development. However, A1 and the dorsoventral eigenmode could diverge as the cortex expands and folds during development, such as the refinement of the Sylvian fissure and Heschl’s gyrus during fetal development (Mallela et al., 2020; Piccirilli et al., 2023). Furthermore, A1 occupies substantially less cortical surface area than V1 or S1 (Glasser et al., 2016; Morosan et al., 2001). This means that functional connectivity maps seeded from A1 will be less likely to show smooth, long-wavelength variations that are captured by the first three non-global modes (Robinson et al., 2003, 2016). In contrast, the larger size of V1 and S1 seeds would more strongly excite the low spatial frequencies represented by the first three eigenmodes (Cruddas et al., 2026; Robinson et al., 2003). Thus, the auditory cortex may be preferentially driven by higher-frequency, short-wavelength eigenmodes or a linear combination of the first three eigenmodes, beyond simply the low-frequency dorsoventral mode alone.

These results suggest that geometry may more strongly constrain the locations and functional organization of visual and somatosensory, rather than auditory, cortices. While many contemporary brain maps and classical studies have identified vision, somatosensation, and audition as roughly co-equal anchors of sensation (Glasser et al., 2016; Luria, 1970; Mesulam, 1998; Smith et al., 2012), other frameworks have minimized the auditory cortex as a dominant pillar of sensory function (Margulies et al., 2016; Uddin et al., 2019, 2023; Yeo et al., 2011). Indeed, fMRI studies often subsume the auditory cortex within pericentral sensorimotor systems rather than distinguishing it as an independent sensory pole (Kong et al., 2025). For example, both the 7-network and 17-network variations of the Yeo-Krienen parcellation derived from resting-state functional connectivity cluster the auditory network within the ventral somatomotor network (Yeo et al., 2011). Our present results show a similar trend; the auditory pole was less pronounced within geometric eigenmode-based sensory integration maps, becoming blurred with the somatosensory pole. Nevertheless, the correspondence between the auditory FC map and dorsoventral mode exceeded chance expectations, indicating that geometry may also play a role in shaping the auditory system.

### The initial spatial layout of primary areas may be guided by geometric constraints

The alignment between geometric eigenmodes and primary sensory fields suggests coarse geometric constraints contribute to unimodal cortical organization. While the dominant rostrocaudal, mediolateral, and dorsoventral eigenmodes are not unique to human cortical geometry, with analogous patterns arising in spheres and other geometries (Gabay & Robinson, 2017; Pang et al., 2023b; Robinson et al., 2003, 2016), this observation may in fact strengthen their relevance to sensory cortical organization. Primary sensory areas mature early in development (Larsen et al., 2023; Sydnor et al., 2021), before extensive cortical folding when the cortex more closely approximates a smooth spherical surface.

Consequently, the low-order geometric modes that characterize simple geometries may provide a parsimonious scaffold for the initial spatial organization of unimodal systems. Nevertheless, the specific spatial embedding of the cortex is necessary to constrain the positioning of modal peaks. While similar low-order modes can arise on any spherical surface, cortical geometry determines how these patterns are oriented relative to sensory landmarks, enabling their correspondence with primary cortical fields

This relationship between simple geometric patterns and sensory field topography appears to be conserved across species (Kaas, 2011; Krubitzer, 2007; Krubitzer & Kaas, 2005; Pang et al., 2025; Xu et al., 2020). The posterior mammalian cortex consistently supports visual function (Butler & Hodos, 2005; Northcutt, 2002), even in non-mammalian species with differently shaped brains or species that rely less upon vision (Catania, 2005), the homolog of V1 occupies an extreme caudal position along the cortical sheet. Similarly, the somatosensory cortex consistently occupies a dorsal and central position across most mammals (Krubitzer et al., 1995).

Furthermore, while the primate auditory cortex lies within the temporal lobe, it is more exposed on the lateral surface of the superior temporal gyrus in non-human primates (Hackett, 2008; Kaas & Hackett, 2000). The deeper position within the lateral sulcus of human A1 suggests a reduced geometric influence along the dorsoventral axis, potentially reflecting closer integration with surrounding association areas to support higher-order language processing across evolution. These cross-species regularities are consistent with the possibility that primary sensory areas occupy locations that exploit conserved geometric features of the cortical sheet. This constraint may result from geometric influences on the spatial expression patterns of cortical patterning genes, which tend to follow the axes corresponding to the first three non-global eigenmodes (Cadwell et al., 2019). The poles of these axes may represent signposts for sensory afferents navigating to their target neurons, triggering the early differentiation of sensory fields from adjacent cortex. This view aligns with evidence that such simple geometric constraints can explain diverse features of regional organization in different mammals (Pang et al., 2025).

### Unimodal-to-transmodal organization requires additional input from successive eigenmodes

Although the first three non-constant geometric eigenmodes broadly described sensory functional connectivity, linear combinations of these three geometric eigenmodes poorly explained the unimodal-to-transmodal axis. One explanation is that the unimodal-to-transmodal gradient depends strongly on transmodal systems such as the default mode network, where long-range connectivity is especially prominent (Oligschläger et al., 2017; Sepulcre et al., 2010). Unlike unimodal primary areas, the default mode network and association cortex contain the majority of long-range connectivity hubs in the brain (Fornito et al., 2025; Oldham & Fornito, 2019), and also exhibit reduced coupling between cortical structure and function relative to unimodal cortex (Valk et al., 2022). Indeed, the removal of these long-range connections causes latent functional connectivity embeddings to resemble a geometric rostrocaudal pattern, rather than the hierarchical unimodal-to-transmodal gradient (Holmes et al., 2026). Modelling hierarchical organization likely requires additional parameters, such as successive geometric eigenmodes or non-geometric features like gene expression and thalamocortical connectivity (Fornito et al., 2025; Müller et al., 2020; Park et al., 2024).

Model divergence in the transmodal cortex may also reflect limitations of sensory angle itself. In low-magnitude regions, corresponding to transmodal association areas, angular estimates are sensitive to small modality-specific fluctuations, which can disproportionately affect angular estimates. Functional connectivity maps contain local variations that are not captured by the first three smooth geometric eigenmodes, but such fine-scale structure can be captured using higher-frequency geometric eigenmodes. Thus, differences between models in transmodal cortex may reflect meaningful fine-scale functional structure, noise in low-signal connectivity estimates, or both.

## Limitations

The present study only conducted analyses at a group, rather than at an individual level. The use of averaged template surfaces can potentially bias results, being less convoluted than individual surfaces (Fischl, 2012). Although the basic patterns of dominant geometric eigenmodes are largely invariant across individuals (Pang et al., 2023a), the specific location of the anchors may vary. The coarser geometry of the averaged template surface may shift modal peaks relative to individual brains (Olsen et al., 2025). This could subsequently affect the correlation magnitudes with sensory functional connectivity, especially with auditory functional connectivity from A1, which is a small region of interest.

Similarly, recent research suggests that cortical geometry is associated with individual variations in functional organization (Alberti et al., 2026; Benn et al., 2025). However, such individual variations predominantly arise in intermediate zones between transmodal and unimodal regions, rather than the unimodal areas that co-anchor geometric eigenmodes and sensory functional connectivity. As such, while the present results would likely remain robust beyond a group-level analysis, future research could utilize sensory tasks to localize A1, V1, and S1, and therefore get individualized mappings between sensory function and geometry .

## Conclusions

Geometric eigenmodes accurately captured sensory integration within primary areas, with visual, somatosensory, and auditory function each aligning with a specific low-frequency geometric eigenmode. Together, the first three non-constant geometric eigenmodes formed a compact space that differentiated unimodal sensory territories. These findings are compatible with the hypothesis that geometry provides a fundamental structural scaffold for the spatial layout of primary areas in the brain.

## Author contributions

Conceptualization: A.H., P.A.R., D.S.M.

Methodology: A.H., W.W., D.S.M.

Software: A.H., W.W.

Investigation: A.H.

Resources: D.S.M.

Data Curation: A.H.

Supervision: D.S.M.

Writing—original draft: A.H.

Writing—review & editing: A.H., W.W., R.A.B., F.A., R.S., J.C.P., A.F., P.A.R., D.S.M.

## Data and materials availability

All data are available in the main text or the supplementary materials, with the raw HCP dataset available online from the original data providers

