## Supplementary Materials for "Cortical geometry constrains the unimodal anchors of sensory integration"

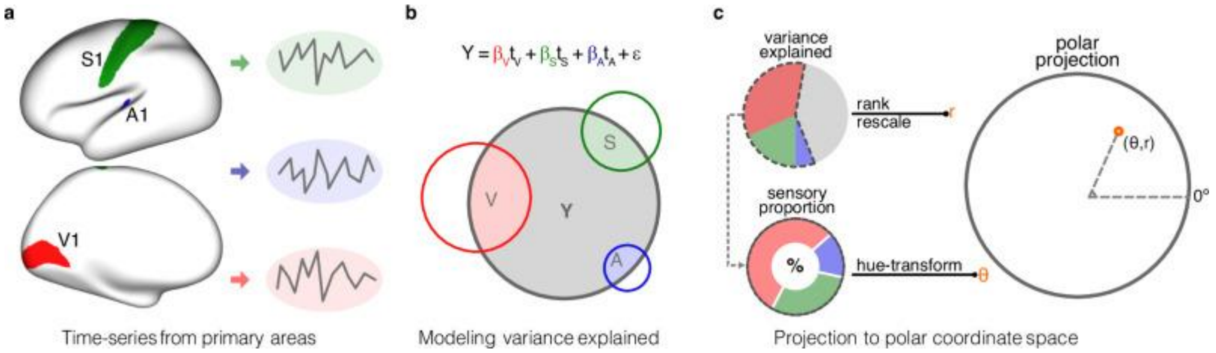


Figure S1: Pipeline to construct the sensory integration model from Wei et al., 2024 under CC BY 4.0. (A) Mean time series was computed separately for V1, S1, and A1 based on the HCPMMP1 parcellation. (B) A non-negative linear model was used to generate sensory components within each vertex by using primary sensory time series as predictors. The lower Venn diagram provides a schematic of the different components within the above equation. (C) Ratios of the variance explained by primary sensory predictors were ranked and rescaled to a range from 0 to 1, representing one dimension of the sensory integration model, and were named magnitude (r). For each vertex, three sensory parameters (βV, βS, and βA) were converted into an angle (θ) using hue transformation, representing the other dimension that indicated the proportional contributions of different sensory modalities.

| Eigengroup | Included Eigenmodes | Wavelength (mm) |
|---|---|---|
| 1 | 1-3 | 297.7 |
| 2 | 4-8 | 171.9 |
| 3 | 9-15 | 121.5 |
| 4 | 16-24 | 94.1 |
| 5 | 25-35 | 76.9 |
| 6 | 36-48 | 65.0 |
| 7 | 49-63 | 56.3 |
| 8 | 64-80 | 49.6 |
| 9 | 81-99 | 44.4 |
| 10 | 100-120 | 40.1 |
| 11 | 121-143 | 36.6 |
| 12 | 144-168 | 33.7 |
| 13 | 169-195 | 31.2 |
| 14 | 196-224 | 29.1 |
| 15 | 225-255 | 27.3 |
| 16 | 256-288 | 25.8 |
| 17 | 289-323 | 24.4 |
| 18 | 324-360 | 23.1 |
| 19 | 361-399 | 22.1 |
| 20 | 400-440 | 21.1 |
| 21 | 441-483 | 20.2 |
| 22 | 484-528 | 19.4 |
| 23 | 529-575 | 18.7 |
| 24 | 576-624 | 18.0 |
| 25 | 625-675 | 17.4 |
| 26 | 676-728 | 16.9 |
| 27 | 729-783 | 16.4 |
| 28 | 784-840 | 15.9 |
| 29 | 841-899 | 15.4 |
| 30 | 900-960 | 14.8 |
| 31 | 961-1023 | 13.4 |

Table S1: Eigengroup membership and wavelengths of the first 1000 non-constant eigenmodes, across 31 eigengroups. Note that the 31st eigengroup contains eigenmodes beyond 1000, yet only the first 1000 eigenmodes were used within the present study. Wavelengths for each eigengroup were calculated following Eq. 4 in the Methods section using a sphere with radius $R_s = 67$ mm.

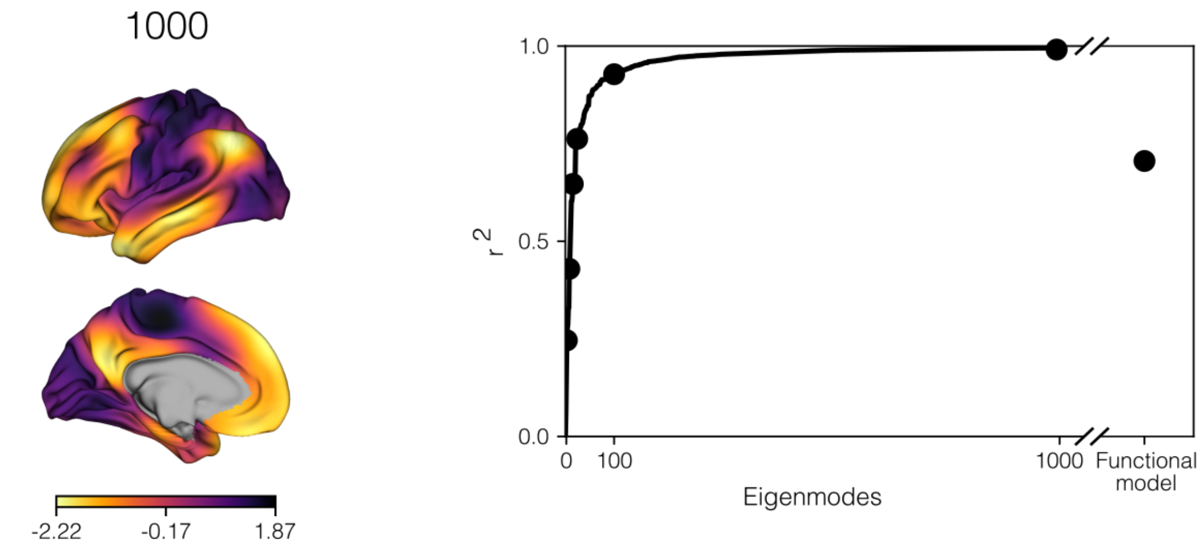


Figure S2: Reproduction of sensory magnitude and the unimodal-to-transmodal hierarchy from an increasing number of eigenmodes, up to a maximum of 1000 modes. Black datapoints within the figure reflect values at 3, 8, 15, 24, 100 eigenmodes (as in Figure 2), alongside an additional value at 1000 eigenmodes, where $r^2 = 0.995$.

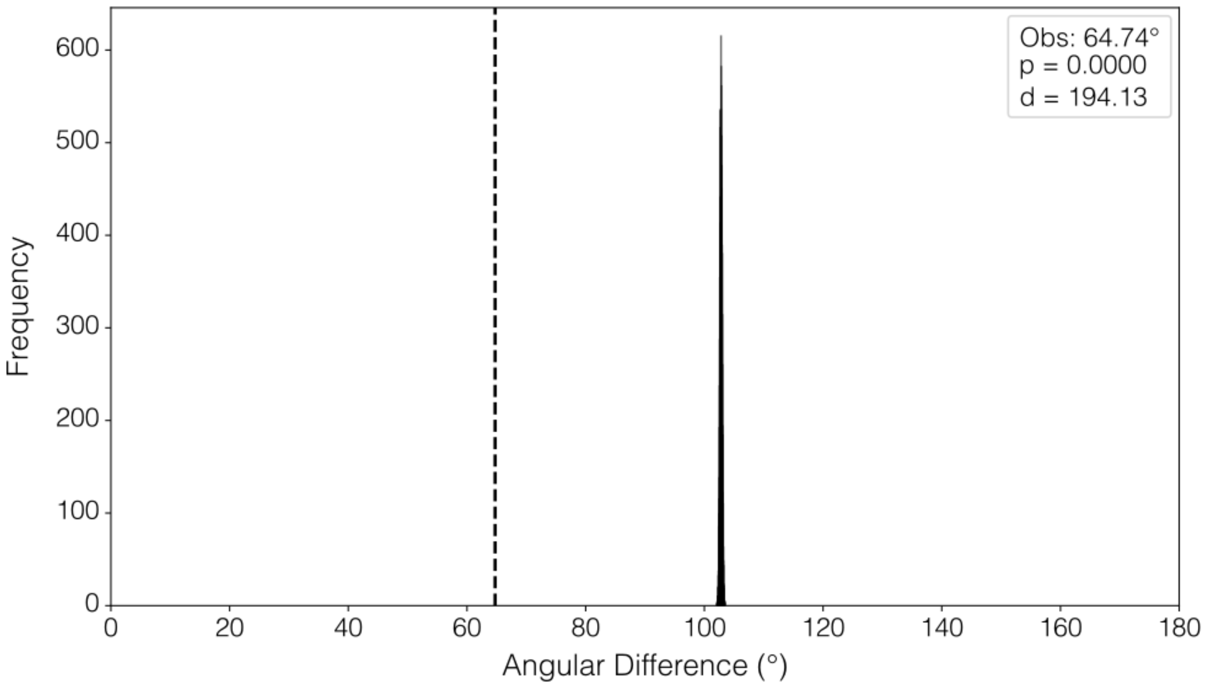


Figure S3: Null distribution from permutation testing of the angular difference between functional and geometric models. The histogram shows the distribution of angular differences obtained from 10,000 random permutations, while the dashed vertical line indicates the observed angular difference (64.74°). The observed value was significantly lower than the null distribution ($p < 0.0001$; Cohen's d = 194.13), indicating greater similarity and fewer differences between models than expected by chance.
